# Genomewide identification of subtelomeric silencing factors in budding yeast

**DOI:** 10.1101/2022.07.20.500793

**Authors:** Alejandro Juárez-Reyes, J. Abraham Avelar-Rivas, Jhonatan A. Hernandez-Valdes, Bo Hua, Sergio E. Campos, James González, Alicia González, Michael Springer, Eugenio Mancera, Alexander DeLuna

## Abstract

Subtelomeric gene silencing is the negative transcriptional regulation of genes located close to telomeres. This phenomenon occurs in a variety of eukaryotes with salient physiological implications, such as cell adherence, virulence, immune-system escape, and aging. The process has been widely studied in the budding yeast *Saccharomyces cerevisiae*, where genes involved in this process have been identified mostly on a gene-by-gene basis. Here, we introduce a quantitative approach to study subtelomeric gene silencing, that couples the classical *URA3* reporter with *GFP* monitoring, amenable to high-throughput flow cytometry analysis. This reporter was integrated into several subtelomeric loci in the genome, where it showed a gradual range of silencing effects. By crossing strains with this dual reporter at the *COS12* and *YFR057W* subtelomeric query loci with gene-deletion mutants, we carried out a genome-wide, comprehensive screen for subtelomeric-silencing factors. The approach was replicable and allowed detection of expression changes caused by previously described silencing factors. We also identified new molecular players affecting this process, most of which are related to functions underlying chromatin conformation. This was the case of *LGE1*, a novel silencing factor herein reported, associated with histone ubiquitination. Our strategy can be readily combined with other reporters and gene perturbation collections, making it a versatile tool to study gene silencing at a genome-wide scale.

## INTRODUCTION

The condensation level of chromatin varies along the genome and impinges on a variety of cellular processes. One of the most important consequences of chromatin compactness is the accessibility of the transcriptional machinery that orchestrates gene expression. In general, highly compacted chromatin regions (heterochromatin) are associated with low transcription rates whereas loosely packed regions (euchromatin) are accessible chromatin sites that are transcriptionally active. In *Saccharomyces cerevisiae*, heterochromatic-like regions are well localized to telomeres, the silent mating type loci, and rDNA repeats, making the budding yeast an excellent model organism to study chromatin conformation. At telomeres, the chromatin condensed state extends to its adjacent regions (subtelomeres) producing transcriptional inactivation or “silencing” of the genes in these loci. This phenomenon has also been termed telomere position effect (TPE) and, overall, it has been associated in different eukaryotic organisms to a variety of traits such as aging (Kaeberlein et al. 1999), cell adherence (Castano et al. 2005; Su et al. 1995), virulence (De Las Penas et al. 2003; Duraisingh et al. 2005; Tham and Zakian 2002; Janzen et al. 2004), along with other features of industrial relevance (Halme et al. 2004; Bauer et al. 2010)

The silenced state at telomeres in *S. cerevisiae* is produced mainly by the SIR complex constituted by Sir4, Sir3 and Sir2 (Aparicio et al. 1991). This complex is recruited to telomere ends by Rap1(Liu and Lustig 1996), which binds specific DNA sequences at the telomere repeats termed silencers. Occupancy of the SIR complex at the subtelomeric regions is propagated inwards, continuously towards the centromere through the action of the histone deacetylase Sir2 (Hoppe et al. 2002). Interestingly, the SIR complex is also one of the multifunctional complexes involved in telomere homeostasis (Kupiec 2014). The deletions of Sir3 and Sir4 both cause shortening of telomeric repeats and mitotic instability of chromosomes (Palladino et al. 1993). TPE is also influenced by gene-dosage balance of telomeric and subtelomeric complex components (Renauld et al. 1993). For instance, Sir3 overexpression causes spreading of silencing over longer distances from the telomere (Hecht et al. 1996). Besides the protein complexes that exert silencing, the size and structure of the telomere tract also influence TPE. It has been observed that short telomeres are associated with diminished TPE (Kyrion et al. 1993) and that telomere folding is also relevant for the maintenance of TPE (de Bruin et al. 2000). In addition, it is known that chromosome context influences silencing levels; regulatory elements at the subtelomeric regions contribute to the intrinsic basal silencing level of each subtelomere (Mondoux and Zakian 2007).

Over 100 genes have been reported to affect Sir-mediated silencing levels at different telomeres in *S. cerevisiae*. These genes were identified mostly on a gene-by-gene basis, usually using the *URA3* reporter gene. The classic assay is based on the experiments that unintendedly led to the original discovery of TPE in yeast (Gottschling et al. 1990); it involves growing a strain carrying the *URA3* gene in a silenced subtelomeric region in the presence of 5-fluoro-orotic acid (5-FOA). In 5-FOA containing media, Ura3 activity produces a toxic metabolic intermediate, causing cell death. Therefore, colony growth can be used as a readout for the intensity of subtelomeric silencing, whereby further genetic modifications with impact on gene silencing result in *URA3* expression and, thus, cell death (Boeke et al. 1984). This semi-quantitative assay has inherent drawbacks, since it has been reported that 5-FOA induces metabolic changes leading to apparent TPE effects in some gene mutants (Rossmann et al. 2011). In addition, the assay is labor intensive and not amenable to testing hundreds or thousands of mutant strains.

In principle, any methodology to measure gene expression such as RT-qPCR or RNA-seq can be employed to assess subtelomeric gene silencing. However, due to labor and cost, most of these methods cannot be readily used in combination with gene-deletion or other available strain collections allowing genome-wide genetic analysis. In this work, we developed a screening approach based on a novel *URA3-GFP* dual reporter integrated into subtelomeric loci to evaluate the effect of non-essential gene knockouts (Giaever et al. 2002) on silencing using high-throughput quantitative flow cytometry. In contrast to other techniques to measure gene expression, flow cytometry is less expensive, suitable for large-scale screenings, and does not require nucleic acid isolation. In addition, gene expression data is obtained at single-cell resolution in live cells, allowing the analysis of not only changes in average expression levels, but also changes in the distribution of gene expression levels across a population. By using this robust and sensitive approach, we reveal variation in gene silencing among different subtelomeric regions of the genome and score genes influencing this phenomenon. Our study provides a large-scale screening approach to pinpoint genes and functions with impact on subtelomeric gene silencing.

## RESULTS

### A dual *URA3-GFP* gene reporter system allowing a quantitative assessment of gene silencing

To screen for genes that influence subtelomeric gene silencing in budding yeast, we constructed a dual-reporter system consisting of a translational fusion of the *URA3* and *GFP* genes under the transcriptional control of the silencing-sensitive *URA3* promoter (Materials and Methods). The *URA3* gene with its native promoter has been widely used to detect gene silencing (Gottschling et al. 1990), but the addition of the *GFP* gene to the construct allows assessing gene silencing by fluorescence microscopy, and, more importantly, by flow cytometry which makes the system amenable to high-throughput screening.

To test the dual-reporter system, we inserted the cassette at two loci that are known to be silenced, the *COS12* and *YFR057W* genes (Mondoux and Zakian 2007; Vega-Palas et al. 2000) at the subtelomeric regions of chromosomes VII (left arm) and VI (right arm), respectively (**Figure 1A**). These two genes display amongst the highest fold increase in expression in a *sir3*Δ mutant (Wyrick et al. 1999), suggesting that the silenced state is mediated by the SIR complex. Furthermore, the telomere where *COS12* resides often localizes to the nuclear periphery, a naturally silencing-promoting nuclear location (Tham et al. 2001). *COS12* belongs to the large family of *COS* subtelomeric genes, a poorly studied set of genes, most of which are the first protein-coding gene next to the conserved core X element of the chromosome. The double reporter was integrated by replacing the entire open reading frames (ORFs) of *COS12* or *YFR057W*, in a strain that lacks the native *URA*3 gene. In this way, silencing of the reporter can be tested in one side by growing the strains in media lacking uracil or containing 5-FOA, and in the other by measuring GFP fluorescence. As a control for non-silenced gene expression, we inserted the dual cassette at the large intergenic region between the *CUP9* and *TRE1* loci in the left arm of chromosome XVI. Parental strains also expressed mCherry from a strong constitutive promoter.

**Figure 1.**
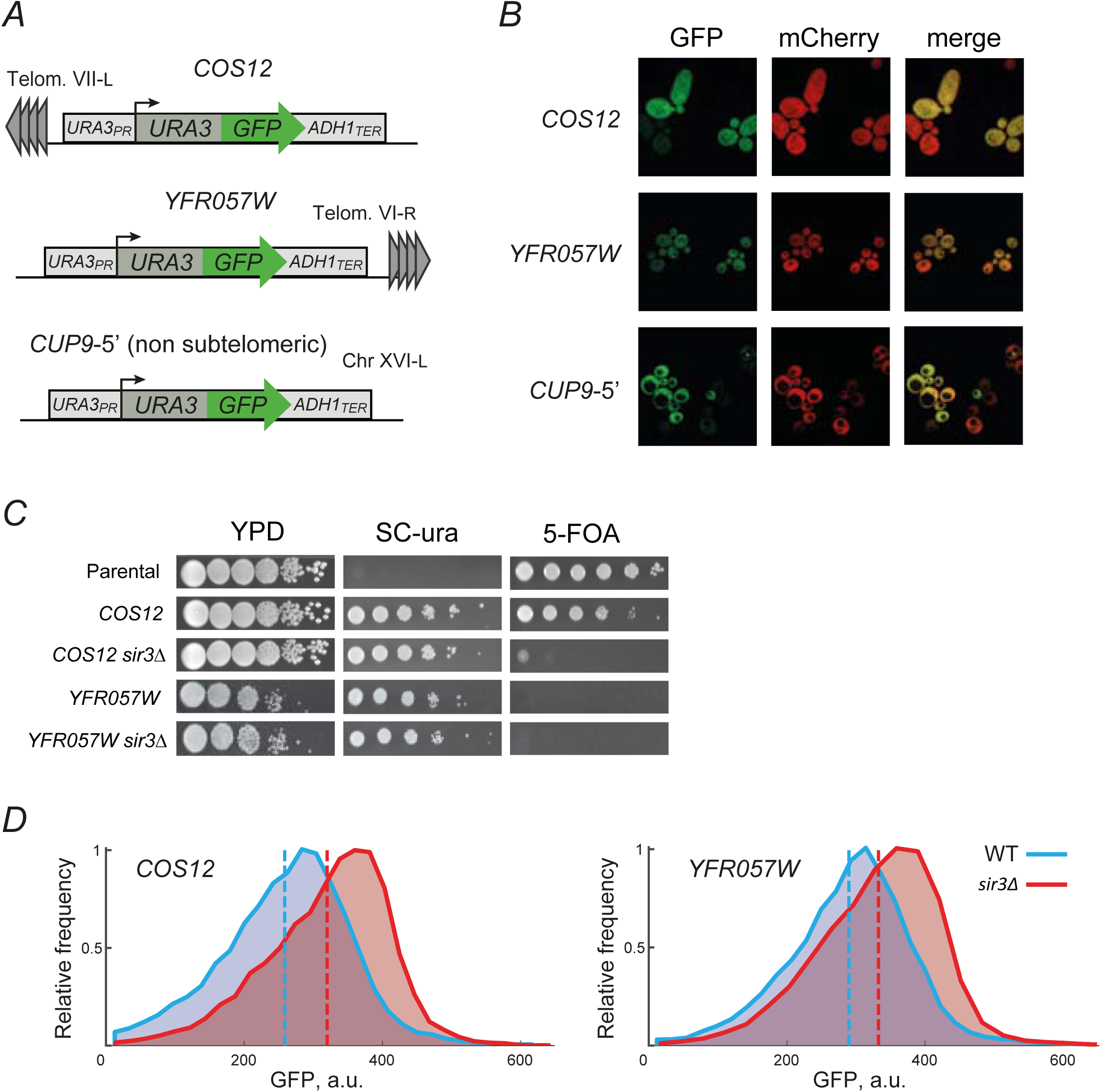
A dual-reporter system to assess gene silencing by flow cytometry. (**A**) Schematic representation of the *URA3-*GFP reporter cassette and its integration at the subtelomeric loci *COS12* and *YFR057W* by replacing the open reading frames, and at the non-subtelomeric *CUP9-5’* intergenic region. (**B**) Confocal fluorescence microscopy images of *S. cerevisiae* cells, bearing the *URA3-GFP* reporter integrated at subtelomeric loci *COS12, YFR057W* and at the not subtelomeric locus *CUP9*. Strains also express mCherry constitutively; GFP and mCherry channels are shown. (**C**) 5-FOA growth assays of the parental strain (*ura3*Δ) and the *URA3-*GFP integrations at *COS12* and *YFR057W* loci in a parental WT or *sir3*Δ background. (**D**) Distribution of GFP signal measured by flow cytometry in the strains where the double reporter is inserted at *COS12* or *YFR057W* in the parental (blue) or *sir3*Δ (red) strain backgrounds. Dashed lines show the mean GFP signal for each cell population.

GFP fluorescence was detected by confocal fluorescence microscopy in cells carrying the reporter in all tested loci, showing that GFP expression from the reporter is functional (**Figure 1B**). We observed reduced GFP signal in yeast cells with the GFP reporter inserted at both subtelomeric loci, especially in *COS12*. It must be noted that such silencing occurs in a variegated manner, namely that GFP signal is very low in some cells while higher in others. Such variegated gene expression is usually observed at silenced loci in yeast [20]. The GFP signal at the non-subtelomeric *CUP9*-5’ locus was also variable, but overall higher. To test the reporting potential of the system based on uracil metabolism, we grew the strains in media lacking uracil or containing 5-FOA (**Figure 1C**). Strains carrying the dual reporter at both subtelomeric *COS12* and *YFR057W* loci were able to grow in medium lacking uracil. However, only the strain with the reported inserted at *COS12* was able to grow in 5-FOA medium. This result confirmed that silencing is incomplete at either locus and is indeed stronger at *COS12*. In fact, growth of the strain with the reporter at *YFR057W* in the presence of 5-FOA was not observed, as if silencing was not occurring at this locus. We also tested the dependency of silencing on the SIR complex by inserting the reporter in a *sir3*Δ strain. As expected, in this background even the strain with the reporter inserted at *COS12* was not able to grow on medium containing 5-FOA, showing that reporter silencing is fully dependent on the integrity of the SIR complex. This observation was quantitatively confirmed by flow cytometry of cells in the late-log growth phases. The mean GFP expression in cells with the reporter inserted at the *COS12* and *YFR057W* loci was higher in the *sir3*Δ compared to the WT background, indicating the SIR-dependency of expression silencing of the *URA3*-GFP reporter by telomere-position effect (**Figure 1D**).

Together, these results show that the dual *URA3-GFP* reporter system allows the measurement of gene silencing level by two independent readouts. First, silencing can be estimated in a semi-quantitative manner using the classical 5-FOA assay based on *URA3* expression and its effect on cell growth, allowing a more direct comparison with previous findings. In addition, GFP fluorescence measurements by flow cytometry provide a quantitative readout that is amenable to high-throughput screening and that is more sensitive to subtle silencing effects, such as that observed at the *YFR057W* subtelomeric locus.

### Subtelomeric regions of *S. cerevisiae* are subject to different levels of gene silencing

The subtelomeric loci *YFR057W* and *COS12* have been thoroughly used to study gene silencing in budding yeast (Mondoux and Zakian 2007; Vega-Palas et al. 2000). Yet, there are 30 other subtelomeric regions in *S. cerevisiae*, many of which remain poorly characterized. To determine the level of subtelomeric gene silencing throughout the genome and to understand whether silencing at *YFR057W* and *COS12* are representative of overall subtelomeric silencing, we integrated the dual *URA3-GFP* reporter at seven other members of the *COS* gene family, each located in the vicinity of different telomeres (see **Table S1** for insertion sites and chromosome features). These genes are not essential and represent, in all but one case, the first gene adjacent to the subtelomeric core X element at the centromere-proximal side. As for *YFR057W* and *COS12*, the reporter was integrated by full replacement of each ORF.

Different reporter expression levels were observed in the subtelomeric-insertions, as inferred from the strain’s capacity to grow on 5-FOA, ranging from full growth of the *COS8* insertion (strongest reporter silencing) to almost no growth in the *COS5* insertion (no silencing) (**Figure 2A**). These *COS8* and *COS5* extreme cases behaved similarly to the parental no-expression and non-subtelomeric unsilenced controls, respectively. In terms of silencing reported by 5-FOA growth, the *COS12* and *YFR057W* insertions were also two extreme cases of strong and undetectable silencing, respectively. Several insertions in the telomere vicinity resulted in little or no apparent silencing in the semiquantitative 5-FOA assay; such was the case of the *COS4, COS2, COS10, YFR057W*, and *COS5* insertions. However, all strains with subtelomeric integrations showed decreased GFP expression compared to the strain with the chromosomal *CUP9*-5’ insertion (**Figure 2B**). The silencing strength determined by GFP expression throughout the subtelomeric loci is like that revealed by growth in the presence of 5-FOA, but at higher quantitative resolution. For instance, there was no growth on 5-FOA of the strain with the reporter at the *YFR057W* locus, which was mostly indistinguishable from the non-telomeric *CUP9*-5’ control insertion; in contrast, the quantitative flow-cytometry assay showed that mean GFP expression was 1.64-fold lower in the *YFR057W* compared to the *CUP9*-5’ insertion. Therefore, these assays show that GFP expression at single-cell level, measured by flow cytometry, resolves slight differences in silencing levels compared to the more qualitative, conventional assay based on *URA3* expression and effect on cell growth.

**Figure 2.**
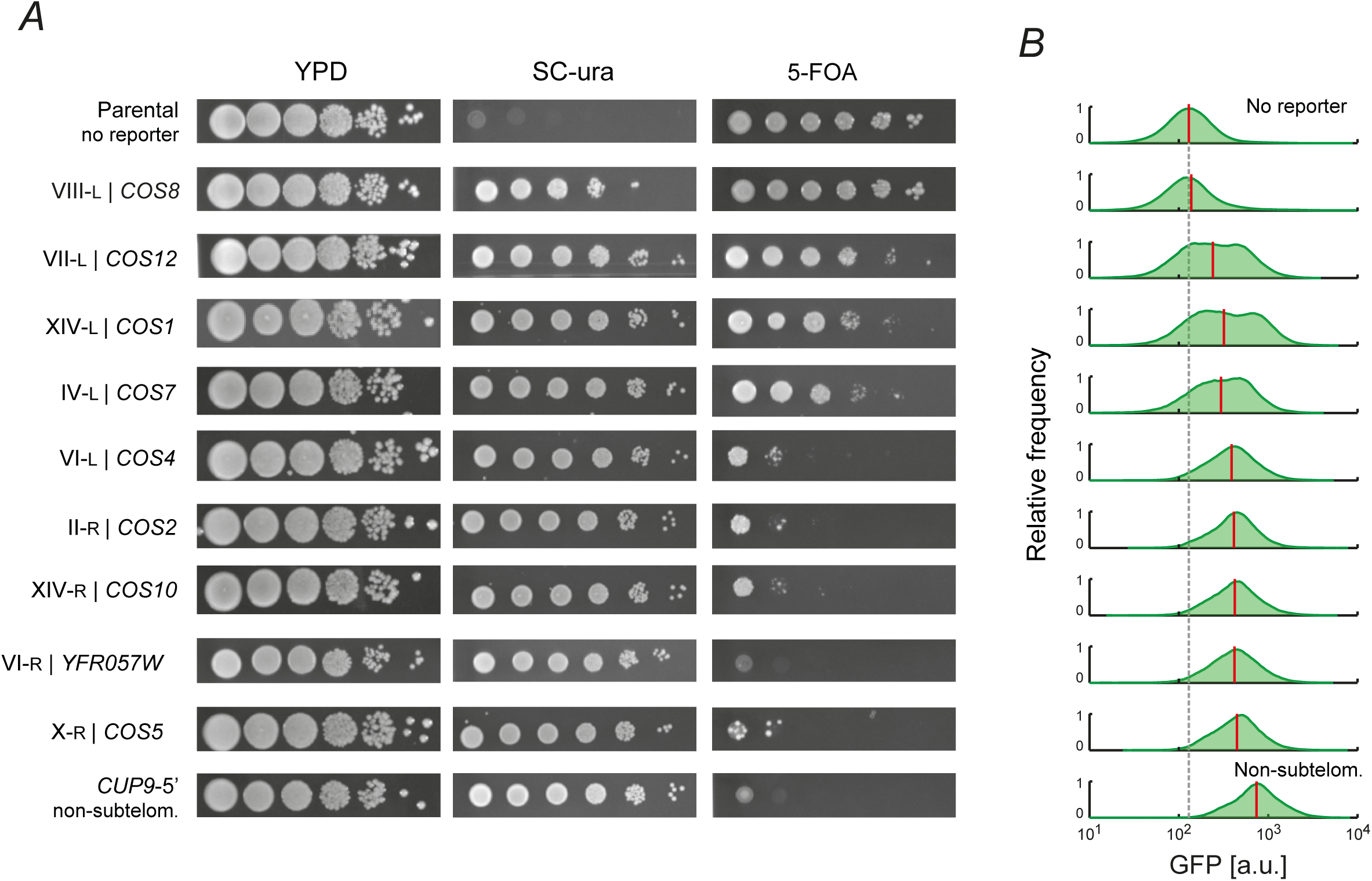
Gene silencing varies across different subtelomeric regions of the yeast genome. (**A**) Strains bearing the *URA3-GFP* dual reporter integrated at the indicated subtelomeric locations by replacing the native ORF were subjected to 5-FOA growth assays. The parental strain (no reporter) is *ura3*Δ. YPD and SC -URA plates were incubated for 48h, while 5-FOA plates were incubated for 72h, all at 30°C. (**B**) Distribution of bulk GFP signal measured by flow cytometry. Strains were grown on liquid SC +20 mg/L uracil and assayed by flow cytometry in the late-log phase. The gray dashed line is the mean GFP background signal of the parental strain, while the red vertical lines indicate the mean GFP signal of each insertion. Integration at the *CUP9-5’* intergenic region was used as a non-subtelomeric reference.

Our results indicate that there is a varying level of gene silencing in the subtelomeric regions of the *S. cerevisiae* genome. A simple explanation for such variation could be the differences in distance of the *COS* genes to the telomere. However, we did not observe such relation of ORF’s ATG distance to the telomere (r = -0.18, *p*>0.05; Pearson) or to the core X element (r = -0.17, *p*>0.05; Pearson) (**Table S1**). Hence, it is likely that other factors of the subtelomeric context contribute to the observed differences in silencing of the same reporter. In this study, we focused on the *COS12* and *YFR057W* loci, which not only have been previously studied at the smaller scale, but also cover the range of silencing strengths of the subtelomeric regions in *S. cerevisiae* as revealed from our results.

### Subtelomeric-silencing factors revealed by genomewide screening

Over 100 genes are known to influence gene silencing at subtelomeric regions in *S. cerevisiae* (**Note S1**). However, a non-biased effort to identify such genes has been missing. To screen for novel genes or pathways that may be involved in gene silencing in a systematic manner, we generated two collections of non-essential gene knockouts bearing the dual *URA3-GFP* reporter at the *COS12* or *YFR057W* loci. These subtelomeric loci are at the extremes of the silencing intensity spectrum (**Figure 2A,B**) and have previously been used to study mechanisms involved in TPE. These collections were generated using a synthetic genetic array (SGA) approach (Tong and Boone 2006), by crossing strains with the integration of the *URA3-GFP* at either loci with a collection of ∼4,500 knockout strains, each one with a non-essential gene replaced by the KanMX cassette (**Figure 3A**). For competitive flow-cytometry analysis, the *URA3-GFP* integrations were done in a strain background constitutively expressing the fluorescent mCherry protein (RFP) in the neutral *HO* locus and an isogenic wild-type strain was labeled with the blue fluorescent mTagBFP2 protein (BFP). The resulting deletion strain collections carry the *URA3-GFP* reporter at *COS12* or *YFR057W*, a KanMX gene replacement and express RFP constitutively.

**Figure 3.**
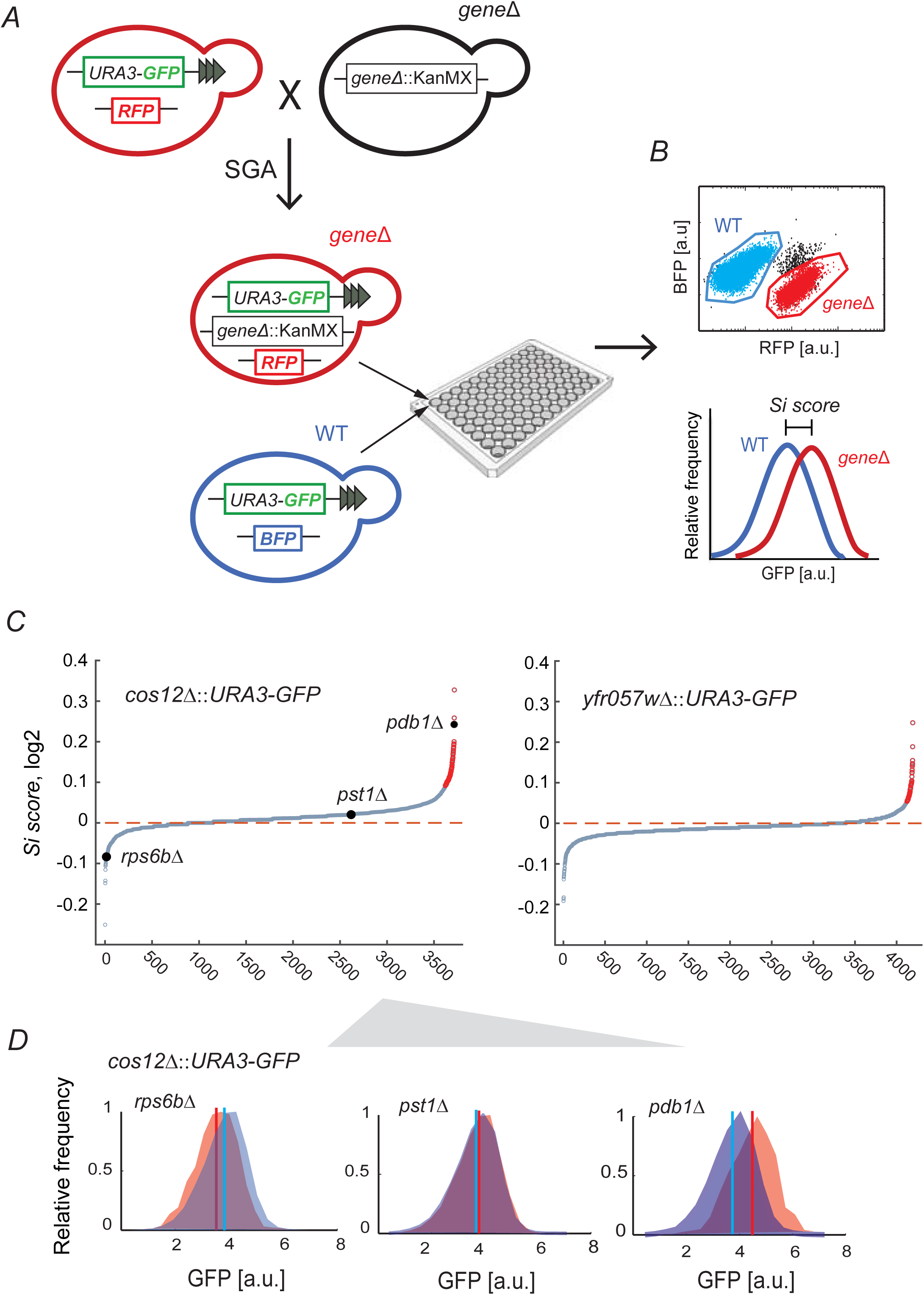
Genome-wide identification of genes affecting subtelomeric gene silencing by high-throughput flow cytometry. (**A, B**) Schematic representation of the screen for subtelomeric gene silencing. (**A**) A large collection of mCherry-expressing knockout mutants harboring the subtelomeric *URA3-GFP* reporter at either the *COS12* or the *YFR057w* locus was generated by SGA (Tong and Boone 2006). For this, a parental strain carrying the reporter at either locus was crossed with a gene deletion collection generated using the KanMX marker (Giaever et al. 2002). (**B**) Each of the resulting mutants was grown in co-culture with a BFP expressing reference strain harboring the subtelomeric *URA3-GFP* reporter at the same locus, with no gene deleted. GFP expression of each pair of mutant RFP and reference BFP strains was measured simultaneously by flow cytometry; separation of the populations was done using their constitutive RFP or BFP signals. The ratio of the average GFP signals of the mutant strain and reference strains was defined as the silencing score (*Si score*). (**C**) Cumulative distribution of *Si score* obtained from screening the gene deletion collection with the reporter at *COS12* (*n=*3,716) and *YFR057W* (n=4,193). Strains that overexpress GFP are marked in red (FDR<10%). (**D**) Distribution of GFP expression of representative strains with distinct Si scores. Mutant strain (red), reference strain (blue); vertical lines are the average GFP signal of each population.

To analyze whether insertion of the *URA3-GFP* reporter affects the local chromatin state in one of the subtelomeric queries, we performed nucleosome-scanning assays (NuSAs) of the *YFR057W* promoter in the wild-type and *yfr057w*Δ*::URA3-GFP* strains. Nucleosome positioning was very similar in the two strains, suggesting that insertion of the reporter had little or no effect on chromatin state in the query strain (Figure S1).

To measure the effect of the deletion of each non-essential gene on subtelomeric silencing in the two query strains, we used high-throughput flow cytometry to measure GFP expression. For increased comparative resolution, each RFP-labeled knockout strain bearing the dual reporter at *COS12* or *YFR057W* was grown in co-culture with the isogenic BFP-labeled wild type, allowing to tell apart the GFP signal of the knockout and wild-type populations in each sample (**Figure 3B**). Typically, between 5000 and 15000 cells were measured from each competitive population. A silencing score (*Si score*) was defined as the ratio of average GFP signals of the mutant and the wild-type reference strains. Based on this metric, we observed that many gene deletions resulted in diminished gene silencing (higher GFP signal, *Si*>1), while others resulted in increased silencing (lower GFP signal, *Si*<1) (**Figure 3C**). Representative GFP expression histograms for knockouts with increased (*rpsb6b*Δ), unaltered (*pst1*Δ), and strong diminished silencing (*pdb1*Δ) are shown in **Figure 3D**. To assess experimental replicability, we screened a fraction of the deletion strains with the *COS12* insertion in two independent experiments, which showed a good rank correlation **(Figure S2**; *r=*0.63, *p<10*^-168^, Spearman).

Using a 10% false-discovery rate, 69 and 55 deletions resulted in decreased gene silencing at the *COS12* and *YFR057W* loci, respectively, while 8 and 25 resulted in increased silencing. There was a trend to more gene deletions having a negative effect on silencing at the *COS12* locus; this trend was less evident at *YFR057W*, which could be associated with the higher basal expression at the later compared to the former locus. Importantly, the *Si score* at both loci are significantly correlated for the 3,677 gene-deletion mutants that were successfully screened in both assays **(Figure 4A**; *r=*0.56, *p<10*^-301^, Spearman).

**Figure 4.**
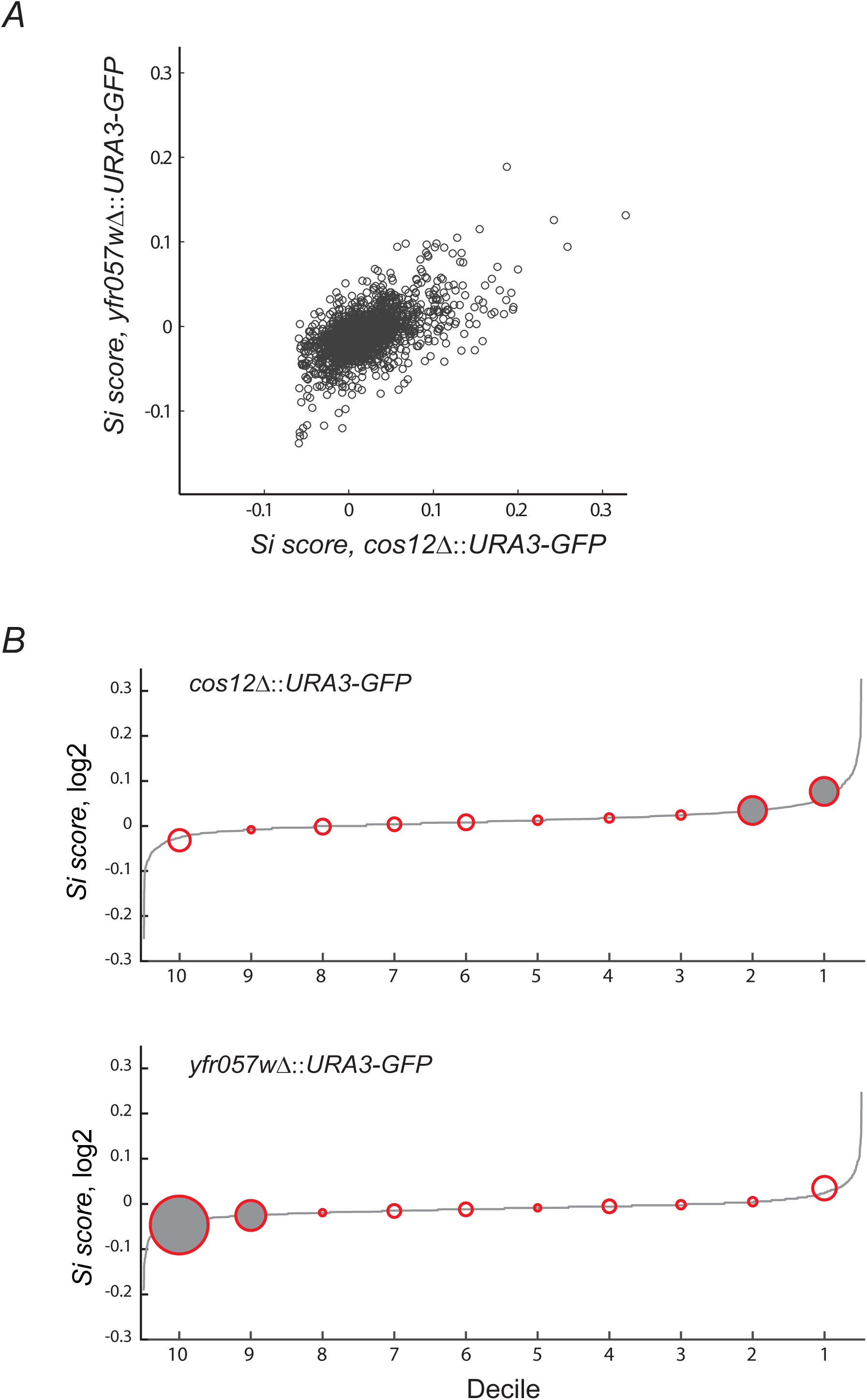
Silencing effects are correlated in both loci and pinpoint previously known genes involved in subtelomeric silencing. (**A**) Comparison between the Si scores obtained at the *COS12* and *YFR057W* loci from the genome-wide silencing screens (*n=*3,677; *r=*0.56, *p*<10^−301^, Spearman). (**B**) Enrichment of genes previously known to be involved in gene silencing at the extreme deciles of the Si score distribution from the screens at *COS12* and *YFR057W*. A list of 132 previously known silencing genes from SGD, from an extensive revision of silencing and from our own curation of the literature (Note S1), was scored at the cumulative distribution of *Si score* divided into deciles. Most of the known silencing genes are at both ends of the distribution. The width of each red circle indicates the number of known genes found at each decile and gray circles denote statistically significant enrichment (*p*<0.05, hypergeometric test).

To assess the quantitative resolution of our approach, we selected a subset of 41 hits above the 10% false discovery rate. These hits included subunits of the main protein complexes identified, 18 mitochondrial genes within the top hits, and individual genes that were not part of an evident complex or functional group. We used the same flow cytometry strategy to measure changes in GFP signal in the *COS12* insertion by performing five technical replicates in competition assays with the BFP-labeled WT reference strain. We used the RFP-labeled *sir3* deletion mutant and the parental WT strain bearing the *URA3-GFP* insertion as positive and negative controls, respectively. Of the re-tested hits, 92.6% showed a significant increase in GFP signal compared to that of the parental reference (**Figure S3**; *p*<0.05, t-test), 87.8% (p<0.01, t-test), 82.9% (p<0.001, t-test). It must be noted that most validated hits showed a modest *Si score*, yet several mutants showed average values above 0.5, including the *sir3*Δ control. Together, these results suggest that screening of changes in expression of the GFP reporter inserted at subtelomeric loci provides a robust, straightforward way to screen for genetic factors involved in gene silencing.

### Silencing effects are consistent with the literature and reproducible between the two readouts of the reporter system

To further validate our genomewide screens, we tested whether previously described silencing genes were overrepresented at the tails of the *Si score* distribution. To this end, we assembled a catalog of 132 genes from the *Saccharomyces* Genome Database (SGD, Gene Ontology Term: chromatin silencing at telomere), an extensive revision of the subject (Mondoux and Zakian; 2006), and our own curation of the literature (**Note S1**). Of the 132 genes, 72 were evaluated in the *COS12* screen and 54% belong to the two higher or lower deciles of the *Si score* distribution, while 54% of the 85 that were measured in the *YFR057W* insertion were in the extreme deciles (**Figure 4B**). The observed enrichments strongly suggest that our large-scale screens revealed genetic factors involved in subtelomeric gene silencing, especially if we consider that the reference catalog includes genes that had been identified in many independent studies, using different methodologies.

Examples of silencing factors confirmed by our screens include genes known to influence telomere length, such as *RIF1* (Hardy et al. 1992), *RIF2* (Wotton and Shore 1997), *YKU70*, and *YKU80* (Williams et al. 2014). The *SPT21* deletion was also part of the top hits in both subtelomeric silencing screens and its knockout mutant has been previously reported to show loss of silencing at subtelomeric positions and altered telomere length (Gatbonton et al. 2006). Spt21 physically interacts with Spt10 (Kurat et al. 2014) and both are required for proper silencing in a native *YFR057W* telomere context (Chang and Winston 2011). In addition, our screens scored other genes related to telomere length (Askree et al. 2004) (*CDC73, RAD50, UPF3)* and telomere capping (Addinall et al. 2008) (*MTC7*). Likewise, different gene knockouts of the elongator complex have been previously reported to diminish silencing of subtelomeric reporters at the VII-L subtelomeric locus, where *COS12* is located (Li et al. 2009). In our work, in both the *COS12* and *YFR057W* screens, deletions of genes of this complex, *ELP2, ELP3*, and *ELP4*, were among the top hits. Another group of genes related to transcriptional regulation obtained at the top positions of the screens were members of the SET3 chromatin remodeling complex (*HOS2, SIF2, SET3*, and *SNT1)*. Interestingly, subunits of the SAS complex showed opposite effects depending on the query locus. For the *YFR057W* locus screen, *sas4*Δ and *sas5*Δ showed a negative effect on silencing, while the subunits Sas2, Sas4, and Sas5 had a positive effect on silencing at the *COS12* locus. These opposite effects were expected since previous studies have shown that components of the SAS complex display locus-dependent opposite silencing effects. In particular, Sas2 activity weakens silencing at a defective *HMR-E* silencer in the *HMR* locus, but promotes it at the *HML* locus and telomeres (Reifsnyder et al. 1996; Ehrenhofer-Murray et al. 1997).

Finally, to validate the results of the screens using the conventional method based on repression of Ura3 activity, we assayed a subset of the top ranked hits for growth on 5-FOA medium. We used the *ura3*Δ knockout strain and the parental *cos12*Δ::*URA3-GFP* insertion as controls. Sixteen out of 21 strains tested (76.2%) showed decreased growth in 5-FOA, suggesting impaired gene silencing at the reporter (**Figure 5A**). As expected, deletion of the *FUR4*-encoded uracil permease results in a mild growth defect, likely due to a direct regulatory effect on the *URA3* promoter and not a telomere-position effect. Among the strains with the strongest silencing defect were mutants of genes known to be involved in subtelomeric gene silencing, such as *yku70*Δ, *yku80*Δ, and *spt21*Δ, which was consistent with their high GFP-signal increase in flow-cytometry validation experiments (Figure S3).

**Figure 5.**
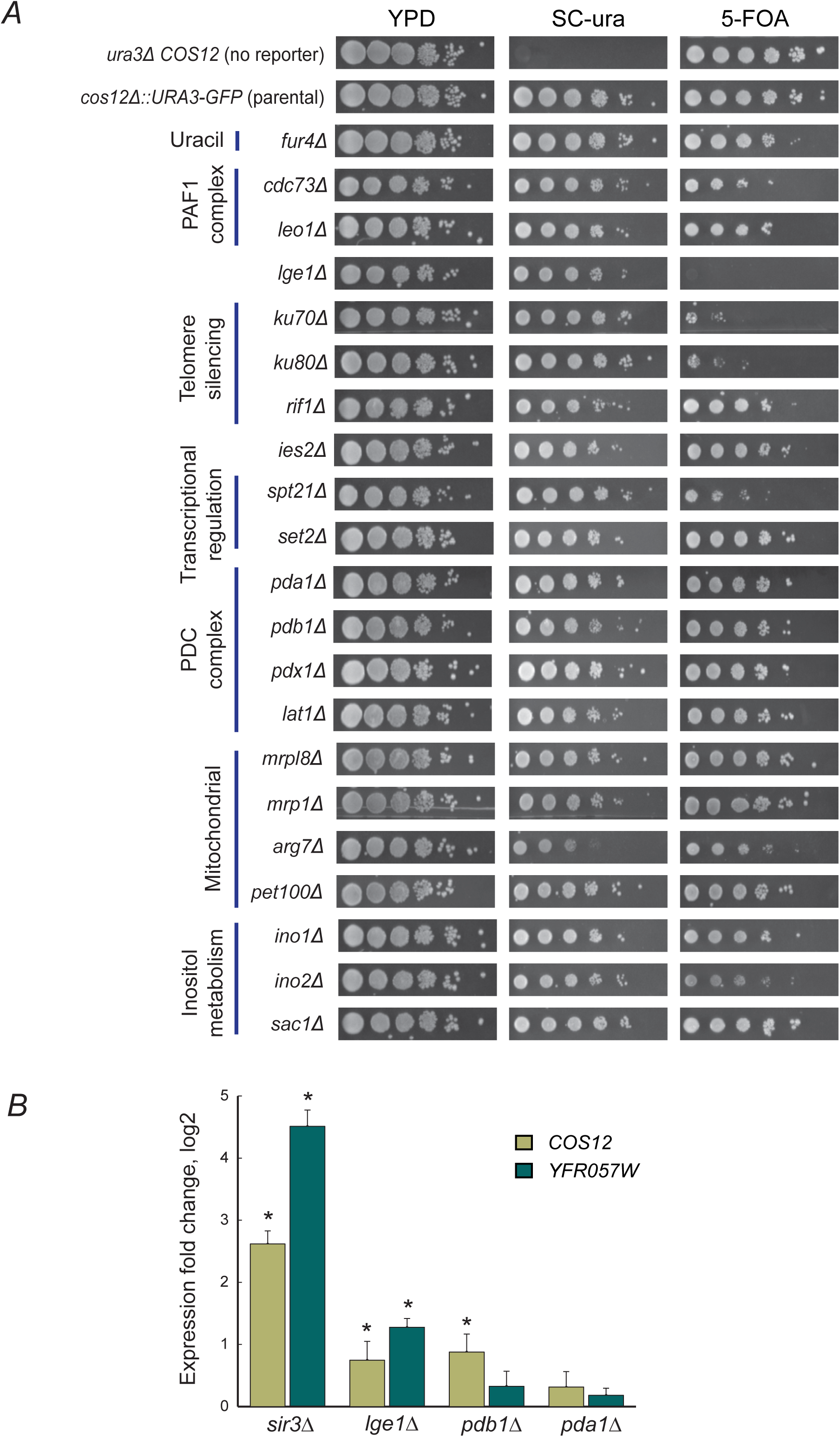
Genes with significantly high *Si score* also show a phenotype in 5-FOA. (**A**) Gene silencing assessment by 5-FOA growth assays at the *COS12* locus of selected strains from overrepresented functional categories or protein complexes. (**B**) Gene expression change of the endogenous *COS12* and *YFR057w* genes in the *sir3*Δ, l*ge1*Δ, *pdb1*Δ, and *pda1*Δ deletion mutants when compared to the parental strain not having the deletion. Gene expression was measured by RT-qPCR in triplicates and expression was internally normalized by *ACT1* expression. Asterisks indicate t-test *p*< 0.01.

### Deletion of *LGE1* results in robust impairment of subtelomeric gene silencing

The *lge1*Δ deletion strain was among the top hits of impaired subtelomeric silencing in both our genome-wide screens. This strain resulted in the most severe 5-FOA growth defect in our validation experiments (**Figure 5A**), which was consistent with the high GFP-signal increase of the *lge1*Δ strain in our highly replicated flow-cytometry experiments (**Figure S3**). Lge1 is involved in H2B ubiquitination mediated by the Rad6/Bre1 complex (Kim et al. 2018), but its precise molecular activity remains unknown. Functionally, Lge1 has been shown to play a role in histone modification and DNA repair, although a direct connection to subtelomeric silencing has not been reported. Given that our reporter system could result in increased expression due to activation of the *URA3* promoter and not a general effect on TPE we used RT-qPCR to test whether the observed effect of *LGE1* impairment was still observed on the native *COS12* and *YFR057W* genes, with no *URA3-GFP* insertion (**Figure 5B**). We observed that both query genes showed a significant two-fold expression increase in the *lge1*Δ compared to the parental strain, indicating that Lge1 activity influences gene silencing independently of effects on the reporter system used. Together, these data confirm that Lge1 is a novel positive subtelomeric silencing factor in budding yeast.

### Expression activation by mitochondrial impairment is not associated to changes in gene expression or nucleosome positioning

We investigated the enrichment of a large set of mitochondrial genes among the mutants with the highest *Si score* in our screens. To this end, we focused on the subunits of the pyruvate dehydrogenase complex (PDC) involved in conversion of pyruvate to acetyl-CoA (*PDB1, PDA1, PDX1*, and *LAT1*). To confirm the effect of PDC subunits on silencing, these knockouts were measured again by flow cytometry at both subtelomeric query loci, which we compared in parallel to subunits of chromatin remodeling complexes well known to affect chromatin structure (Sir3) and were hits in our screens (Set2 and Ies2). All PDC knockouts showed significant *Si score* differences when compared to the WT strain (**Figure 6A**; *p*<10^−2^ and *p*<10^−3^ for *YFR057W* and *COS12* insertions, respectively). Most mutants of the PDC subunits showed stronger effects on silencing of the dual reporter than the mutants of chromatin remodeling complexes that were used as a reference.

**Figure 6.**
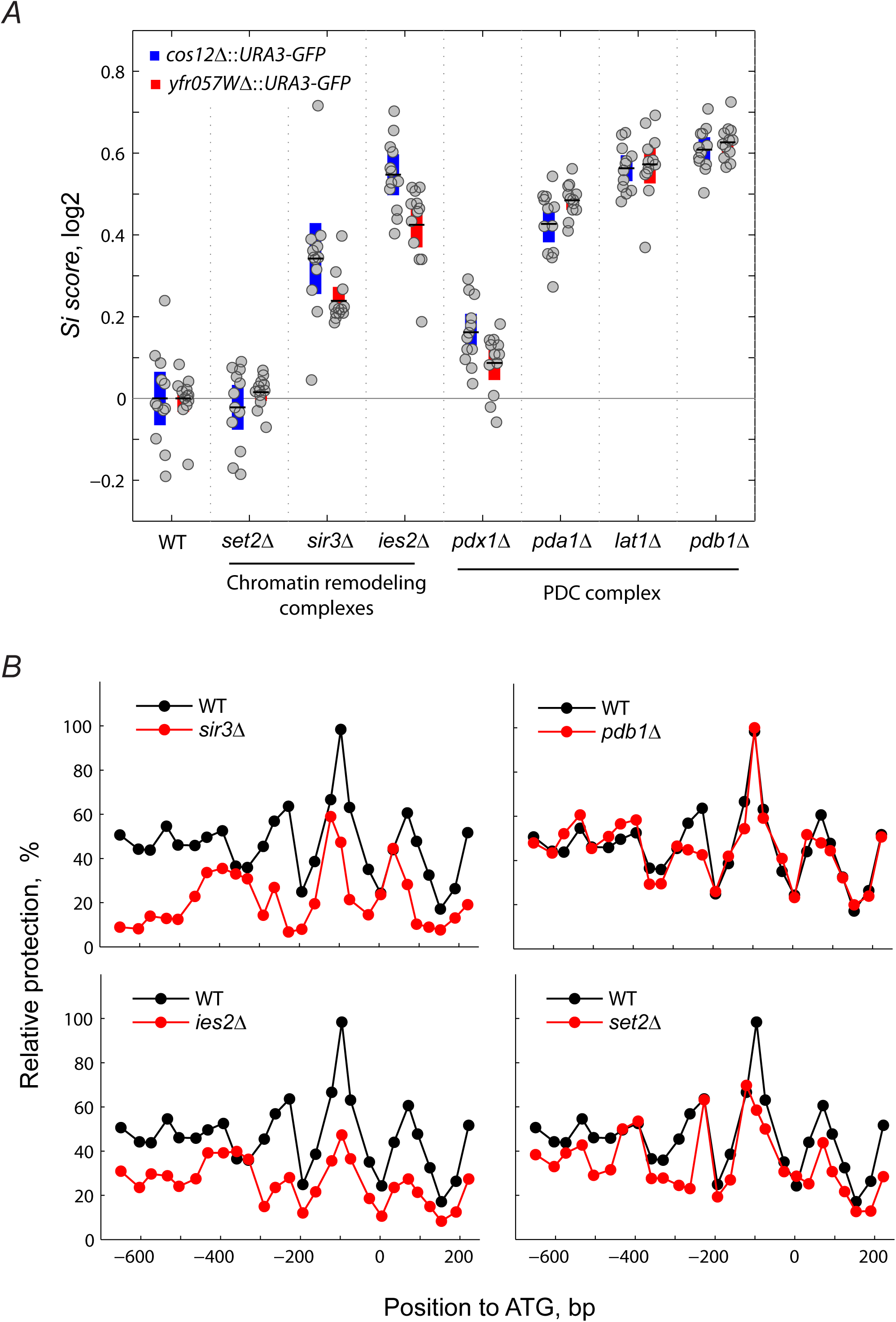
No evidence for a direct role for mitochondrial function in subtelomeric gene silencing. **(A)** Comparison of *Si scores* of knockouts of genes that code for the PDC complex subunits (*pdx1*Δ, *pda1*Δ, *pdb1*Δ, and *lat1*Δ) and chromatin remodeling factors (*set2*Δ, *sir3*Δ, and *ies2*Δ) measured by flow cytometry. The log_2_ *Si score* of the WT was normalized to zero for each locus. (**B**) Nucleosome positioning at the*YFR057W* promoter in the *sir3*Δ, *pdb1*Δ, *ies2*Δ, and *set2*Δ mutant strains. NuSAs were performed on strains that do not have the double reporter by growing them in SC medium containing uracil (20 mg/L) at 30°C and harvested at late-log phase (see Materials and Methods). Relative protection was calculated as a ratio using the *VCX1* gene as a reference since a well-positioned nucleosome is found at the +250 bp position of this ORF. For each primer pair, the midpoint of the PCR fragment is shown as a solid dot and overall, they amplify from around -650 to +222 bp of the *YFR057W* locus. The coordinates are given relative to the ATG (+1).

We tested whether the effects on silencing observed in the mutants of the PDC subunits were due to chromatin changes at the nucleosome level, which are expected in *bona fide* TPE. We carried out nucleosome-scanning assays (NuSA) at the *YFR057W* promoter and the *URA3-GFP* insertion sequences in the *set2*Δ strain and, as a reference, mutants of subunits of chromatin remodeling complexes. In each NuSA assay, nucleosome positioning was compared to the parental strain. Deletion of *PDB1* did not influence nucleosome occupancy at the promoter; nucleosome distribution was very similar to the parental strain. In contrast, the absence of *IES*2, *SIR3*, and *SET2* results in a reduction in nucleosome positioning over the entire promoter region, as expected, even at the well-positioned nucleosome at the -96 bp position (**Figure 6B**). In agreement with its relatively lower *Si score*, the deletion of *SET2* showed the weakest effect on nucleosome distribution at the *YFR057W* query region. Noteworthy, the effects of mutants of PDC subunits *pdb1Δ* and *pda1Δ* on native *YFR057W* expression were also evaluated by RT-qPCR, showing no significant effect on expression (**Figure 5B**). Together, these results suggest that, at least for the top pyruvate dehydrogenase complex hits, the effects of mitochondrial function on gene expression depend on the reporter system used, either due to a higher basal expression or a direct activation effect of the *URA3* promoter.

### A global picture of subtelomeric silencing in yeast

Our genetic screens provide an opportunity to revise the general cellular and molecular functions contributing to subtelomeric silencing, using results from an unbiased genetic dataset. To this end, we used a functional analysis based on Cohen’s *kappa* (Huang da et al. 2009), as previously described (Campos et al. 2018). This analysis evaluates the relationship between gene-pairs by establishing the overall agreement between a set of associated evaluators. Here, evaluators included Gene Ontology (GO) and phenotypic terms that have been previously ascribed to the genes of interest, as reported in SGD. We tested a set of 266 genes (**Table S3**) including 141 hits from our two screens (top and bottom *Si score* rank, FDR <10%) and 125 from the silencing-factors reported in the TPE literature (**Note S1**).

Using the *kappa*-based functional analysis, we identified 11 clusters of genes, each composed of three to a dozen of genes (**Figure 7**). The main cellular processes associated to the clusters were histone and chromatin modification and telomere maintenance. Two high-scoring hits from the screen, *YKU70* and *YKU80*, clustered together with *RRM3* forming a cluster related to telomere maintenance, with known roles on subtelomeric silencing. Most of the observed clusters included genes related to different categories that impact chromatin structure or function. These included genes with roles on nucleosome positioning or remodeling (*ISW2* and *CHD1*) that were connected to the *FUN30* and *INO80* genes. The latter two genes have been previously reported to participate in chromatin silencing. Another cluster included *CDC73, LEO1*, and *RTF1* which all are part of the multifunctional Paf1 complex involved in RNA polymerase II transcriptional elongation, RNA processing, and histone modification during elongation. Interestingly, the novel silencing factor *LGE1* was part of two chromatin-related clusters, linked to histone deacetylases, histone methyltransferases involved in chromatin silencing at telomeres, and other chromatin remodeling complexes. This finding is consistent with the function of Lge1 as a histone H2B-ubiquitination cofactor, and it suggests a possible mechanism through which Lge1 impacts subtelomeric silencing. Our screens also revealed a cluster of several genes involved in ribosomal function and another of uncharacterized ORFs. Further validation is needed to confirm the role of these genes in subtelomeric silencing.

**Figure 7.**
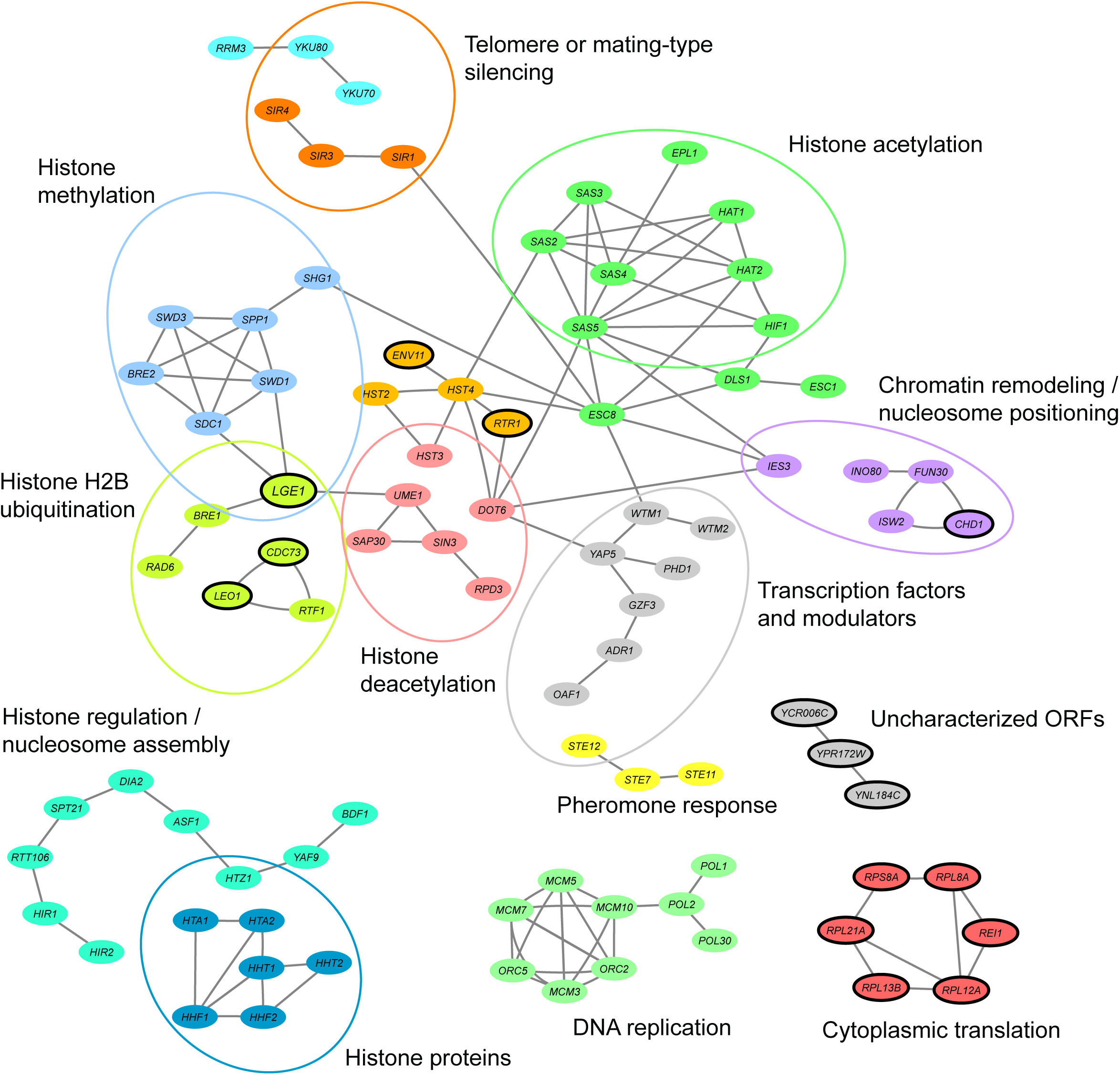
A functional view of subtelomeric gene silencing. Functional annotation of top-ranked *Si score* genes and genes previously known to influence gene silencing by *kappa*-based functional analysis. Functional clusters are represented as a network where genes are oval nodes that are connected by edges when kappa value indicates functional agreement between genes (k>0.35). Colors are used to specify the different clusters within the network. A large cluster of genes with mitochondrial function is not shown, given that effects of mitochondrial function on gene expression do not seem to be due to subtelomeric silencing, but rather to the reporter system used. Ovals with a solid outline indicate genes from the top FDR 10% of the silencing screens, with no previous report on TPE.

## DISCUSSION

Subtelomeric loci are exceptional genomic locations to study gene silencing and its effect on physiological functions. In budding yeast, the model organism where TPE is better understood, this process has been studied genetically factor by factor. Here, we developed a quantitative approach to facilitate the identification of genes that play a role in subtelomeric silencing, by using a double *URA3-GFP* reporter gene system coupled with high-throughput flow cytometry. Our method proved to be more sensitive than classical 5-FOA assays, allowing the detection of subtle differences in gene silencing across a variety of subtelomeric loci in the yeast genome. Telomere length is a known determinant of subtelomeric gene silencing, and we did observe changes in gene expression in mutants altered in telomere length and maintenance. However, distance from the insertion sites of the reporter to the telomere was not correlated with silencing levels, at least within the range of distances that we assessed in nine different insertion sites. Further research will be needed to determine which other factors, such as telomere structure, contribute to the silencing variation that we observed in the different subtelomeric regions of the yeast genome.

We show that the measurement of GFP expression by flow cytometry is a more sensitive readout than the growth on medium containing 5-FOA, even though the results of the two assays were in general agreement, as did silencing levels of a subset of genes that were measured repeatedly by flow cytometry. Furthermore, the top hits identified by our screens are enriched in genes previously known to affect gene silencing. Clear examples are those known to affect telomere length and members of the SET, SAS, Ku70/80 and PAF1 complexes. These results indicate that our approach is robust and amenable to large scale analysis of gene silencing.

Most of the functional categories associated with silencing are related to chromatin conformation and modification, in one way or another. Among these genes, *LGE1* had not been directly associated with subtelomeric gene silencing, but its knockout is one of the mutants that showed the strongest effect in our screens and in the validation experiments. One possible explanation for the role of Lge1 in subtelomeric silencing may be its connection with Set1 and Dot1 (Wood et al. 2003). It is known that monoubiquitination of H2B mediated by Rad6-Bre1-Lge1 is a prerequisite to the H3K4 and H3K79 methylation produced by Set1/COMPASS and Dot1, respectively (Wood et al. 2003). Lysine methylation of H3K79 by Dot1 has been shown to be important during transcriptional elongation by the Paf1 complex and to regulate telomeric silencing (Ng et al. 2002). Thus, it is possible that the loss of Dot1 and Set1 dependent methylation in a *lge1* knockout could affect silencing by disrupting the ability of Sir2 and Sir3 to form heterochromatin. An alternative route, although less clear, could be through the association of Lge1 with the DNA repair protein Ku70. The mutants of these genes show synthetic lethality at high temperature (L. 2008), and Ku70 also shows a synthetic lethal interaction with Set1, the histone methyltransferase that is central for subtelomeric silencing.

Unexpectedly, a large set of genes with mitochondrial function were enriched in our screens. The strongest effect was observed in the mutants of the pyruvate decarboxylase complex, which showed clear and robust increases in subtelomeric GFP expression and impaired growth in 5-FOA. However, further direct expression measurements of the native promoters by RT-qPCR and nucleosome-position analysis of some of the mitochondrial mutants suggested that the effect on silencing is specific to the reporter system used (**Figure 5B** and **Figure 6B**). One possible explanation is that deletion of the mitochondrial genes is specifically interacting with pathways that affect expression from the *URA3* promoter. This interpretation is indeed the case for genes detected by our screens and that are involved in pathways related to the availability of uracil, e.g. the plasma membrane uracil permease Fur4 and the uracil biosynthetic genes *URA5* and *URA1*. Although further work is required to understand the exact connection between mitochondrial genes and the silencing effects observed in our reporter system, our results raise a word of caution for the use of the *URA3* gene for assessing gene silencing, which is routinely done with the use of the 5-FOA growth assay. In future studies it would be very informative to substitute the *URA3* promoter with other promoters in a double reporter assay.

Our screens did not include mutants of genes that are essential or those resulting in sterile strains since they are not amenable to SGA. This is relevant given that some essential genes such as *RAP1* (Kyrion et al. 1993) and *ABF1* (Pryde and Louis 1999) are known to have strong roles in subtelomeric silencing. Similarly, *SIR2* and *SIR3*, whose deletion cause sterility, are main players of subtelomeric silencing. In this work we generated some of the reference strains by direct PCR-based transformation, but essentiality and sterility limitations could be overcome by using strain collections of conditional mutants.

Most subtelomeric genes identified in previous studies were those with strong telomere-position effects. Our work shows that there are many other genes that have subtler effects and that are more readily detected by sensitive, quantitative methodologies. The flow-cytometry based approach presented here also allows obtaining single-cell expression data to identify variegation trends in large populations. Besides being quantitative and amenable to high-throughput screening, our strategy is quite versatile since different promoters and fluorophores can be combined with the many gene deletion collections that are available for budding yeast. We anticipate that using our approach with other promoters and reporters will allow overcoming the caveats detected in our screens, revisit previous studies, and understand novel molecular mechanisms of subtelomeric gene silencing in budding yeast and other organisms.

## MATERIAL AND METHODS

### Strains and strain construction

All strains used in this study are listed in **Table S4**. Knockout strains are from the yeast deletion collection *xxx*Δ::*KANMX4* in the BY4741 background (Giaever et al. 2002). The Y8205 parental mCherry (SCA52) and mTagBFP2 (Subach et al. 2011) (SCA89) fluorescent strains were generated by integrations of fluorescent-NAT cassettes at the HO locus by homologous recombination. Fluorescent-NAT cassettes were constructed on a pFA6 (addgene) based plasmid. All the primers used for the construction of the strains are listed on (**Table S5**). The *URA3-*GFP reporter was PCR amplified from pAJ69 and integrated at subtelomeric loci by homologous recombination using primers sharing 40bp identity with subtelomeric regions. This reporter was integrated into parental strain mCherry (SCA52) and mTagBFP2 (SCA89). Using this methodology, we replaced several subtelomeric genes. The PCR primers in all cases were designed to replace entirely the subtelomeric ORFs of selected genes. At chromosomal internal locus *CUP9* integration occurs at the 5’
s intergenic region leaving intact ORFs. The construction of the library of mutants to study silencing at different loci was based on synthetic genetic array methodology (Tong and Boone 2006). The *sir3*Δ strains for each locus was generated by homologous recombination over the parental strains. Not all crosses and further screening and data acquisition were successful and therefore the final data sets consisted of 3,716 knockouts for the *COS12* locus and 4,193 for the *YFR057* locus.

### Plasmid construction

We constructed plasmid pAJ69 which consists of a translational fusion of *URA3* and GFP genes under control of a minimal *URA3* promoter (216 bp) and *ADH1* terminator. *URA3* gene and promoter were amplified from pRS416 Stratagene and GFP gene and *ADH1* terminator were amplified from pFA6a-GFP (S65T)-His3MX6 (Huh et al. 2003). These PCR fragments were fused by double joint PCR and cloned in pUC19 *EcoRI-HindIII* sites. *URA3* and GFP genes were cloned in frame using a 27 bp linker (primers in **Table S5**).

### Growth conditions for flow cytometry GFP measurements

For large-scale screenings at *COS12* and *YFR057W* loci, the reference BFP strains SCA93 and SCA91 and plates of the respective subtelomeric reporter knockout collection were grown overnight on YPD on 96-well plates at 30°C without shaking. Each pair of reference-mutants were then pinned inoculated in 160 µl SC medium with 20 mg/l uracil in 96-well microtiter plates. Strains were grown at 30°C, 1000rpm for 14-17 hours (7-9 cell generations, OD_600nm_ in microtiter plate reader was 0.4 to 0.6. Cells were treated with 20 µl TE 2X and immediately measured at flow cytometer.

### Flow Cytometry: Instrumentation, acquisition, and data Analysis

Large scale flow cytometry was performed on a Stratedigm S1000EX cytometer. mTagBFP2 was excited with a violet laser (405nm), and fluorescence was collected through a 445/60 band-pass filter. GFP was excited with a blue laser (488nm), and fluorescence was collected through a 530/30 band-pass filter. mCherry was excited with a yellow laser (561nm), and fluorescence was collected through a 615/30 band-pass filter. Each co-culture reference-mutant was set on each well of the plates. The flow cytometer was set to measure 15, 000 events or to stop acquiring after 50 seconds on each well. As a mean for each mutant and reference pair there were more than 10,000 cells counted. The BD FACSCalibur™ and BD LSR Fortessa X-20™ were used for validation in smaller-scale experiments. For BD LSR Fortessa X-20 validation experiments, mTagBFP2 was excited with a violet laser (405nm), and fluorescence was collected through a 450/50 band-pass filter. GFP was excited with a blue laser (488nm), and fluorescence was collected through a 505LP emission filter and a 525/50 band-pass filter. mCherry was excited with a yellow-green laser (561nm), and fluorescence was collected through a 600LP emission filter and a 610/20 band-pass filter. For hit validation of genes of *COS12* screening the flow cytometer was set to measure 30, 000 events or to stop acquiring after 30 seconds on each well. All data analyses and plots were performed with custom scripts on MATLAB.

### Confocal microscopy

Cells were grown on YPD at 30°C to late exponential phase (OD_600nm_ = 0.6-0.9) and then collected and washed thrice with 1 ml PBS 1X (NaCl 8.0 g/L, KCl 0.2 g/L, Na_2_HPO_4_ 1.44 g/L, KH_2_PO_4_ 0.24 g/L), paraformaldehyde 4% fixed, washed again and resuspended in sorbitol 1M solution. Cells were visualized in a LSM800 Zeiss confocal microscope, using 40X or 63X objectives. GFP was excited with a 488 nm laser, and mCherry with a 561 nm laser, and fluorescence was captured using standard parameters and two different channels using filters SP620nm and LBF640.

### 5-FOA growth assays

Strains were grown in YPD medium to stationary phase at 30°C and 200 rpm. The cultures were adjusted to an optical density of 1 at 600nm with sterile water and then 10-fold serial dilutions were made in 96-well plates. A total of 5 µl of each dilution was spotted onto YPD, SC-ura and 5-FOA agar plates and incubated at 30°C for 48h for YPD and SC-ura agar plates and 72h for 5-FOA agar plates and then photographed.

### Nucleosome scanning Assay (NuSA)

Nucleosome scanning experiments were performed adapting the method described previously (Infante *et al*., 2012). The *his3*Δ *S. cerevisiae* considered WT (native and system) and the pertinent mutants were grown to late exponential growth phase (45 mL of an O.D_600_ = 0.8 to 1.0). Cells were treated with formaldehyde (1% final concentration) for 20 min at 37 °C and then glycine (125 mM final concentration) for 5 min at 37 °C. Formaldehyde-treated cells were harvested by centrifugation, washed with Tris-buffered saline, and then incubated in Buffer Z2 (1M Sorbitol, 50 mM Tris-Cl at pH 7.4, 10 mM β-mercaptoethanol) containing 2.5 mg of zymolase 20T for 20 min at 30 °C on rocker platform. Spheroplast were pelleted by centrifugation at 3000X *g* and resuspended in 1.5 mL of NPS buffer (0.5 mM Spermidine, 0.075% NP-40, 50 mM NaCl, 10 mM Tris pH 7.4, 5 mM MgCl_2_, 1 mM CaCl_2_, 1 mM β-mercaptoethanol). Samples were divided into three 500 µL aliquots that were then digested with 22.5 U of MNase (Nuclease S7 from Roche) at 50 min at 37 °C. Digestions were stopped with 12 µl of Stop buffer (50 mM EDTA and 1% SDS) and were treated with 100 µg of proteinase K at 65 °C overnight. DNA was extracted twice by phenol/chloroform and precipitated with 20 µL of 5 M NaCl and equal volume of isopropanol for 1 h at -20 °C. Precipitates were resuspended in 40 µL of TE and incubated with 20 µg RNase A for 1 h at 37 °C. DNA digestions were separated by gel electrophoresis from a 1.5 % agarose gel. Monosomal bands were cut and purified by Wizard SV Gel Clean-Up System Kit (Promega, REF A9282). DNA samples were diluted 1:30 and used in quantitative polymerase chain reactions (qPCR) using primers listed in (Table S5) to quantify the relative MNase protection of *YFR057W* locus template. qPCR analysis was performed using a Corbett Life Science Rotor Gene 6000 machine. The detection dye used was SYBR Green (2× KAPA SYBR FAST qBioline and Platinum SYBR Green from Invitrogen). Real-time PCR was carried out as follows: 94° for 2 min (1 cycle), 94° for 15 sec, 58° for 20 sec, and 72° for 20 sec (30 cycles). Relative protection was calculated as a ratio to the control *VCX1* (*YDL128W*) template found within a well-positioned nucleosome in +250 bp of the ORFs. The PCR primers amplify from around -650 to +222 bp of *YFR057W* locus whose coordinates are given relative to the ATG (+1).

### *Kappa*-based functional analysis

Gene Ontology (GO) and phenotype terms were downloaded from the *Saccharomyces* Genome Database (SGD, last updated October 2019) to build two *m* by *n* matrices, where *m* is the number of analyzed genes 266 and *n* is the number of GO and phenotypic terms (2,234). Each term evaluates the overall agreement between gene-pairs to calculate Cohen’s *kappa* 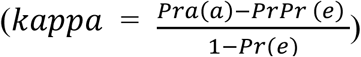. Where *Pra*(*a*) is the number of GO and phenotypic terms in which each gene-pair shares an agreement, divided by the total number of terms downloaded from SGD, and *Pr*(*e*) is the hypothetical probability for each member of the gene-pair to be associated by chance. Then a matrix of *m* by *m* genes representing the agreement as a *kappa* value between each gene-pair was built. Gene-pairs with a *kappa*>0.35 were considered as functionally associated, values above this threshold represent the top 5% *kappa* values, this threshold has also been used in previous reports for large datasets (Campos et al. 2018: Huang *et al*. 2009). In a first step, only gene-pairs are associated, these pairs were then used as cluster seeds to form larger groups of genes with subsequent iterations of the analysis, where clusters sharing over 50% of its members were merged. Later, the clusters were named by manually inspecting for enriched functions or by using GO term finder tool (version 0.86) at SGD. The algorithm for *kappa* analysis was written on Matlab. Cytoscape was used to create a network where associated genes displayed kappa agreement above the threshold (*kappa*>0.35). Analyzed genes are listed in **Table S3**.

### Gene expression by RT-qPCR

Expression of the two subtelomeric genes *YFR057W* and *COS12* was evaluated by RT-qPCR analysis on the WT strain and some deletion strains identified in the screening. In addition, the *sir3*Δ was included as a strain with strong defects in silencing for comparison purposes. For RNA extraction RiboPure-yeast kit (Ambion® by life technologies™) was used. Cells were grown on YPD at 30°C until 0.6 OD_600nm_, then were harvested and put on ice. spectrophotometer. Total RNA extraction was carried out for each strain and cDNA was obtained by triplicate for each RNA extraction following manufacturer’s instructions. RNA integrity was assayed by gel electrophoresis and quantified in NanoDrop ND-1000. cDNA was obtained of 2µg of total RNA using SuperScript™ III Reverse transcriptase and quantified again in NanoDrop. RT-qPCR was performed on StepOne™ Real-Time PCR system (Applied Biosystems), for 40 cycles using power SYBR^®^ Green PCR Master Mix (Applied Biosystems) and primers listed in **Table S5** with a T_a_ = 60°C. A ∆Ct was normalized to *ACT1* for each *COS12* or *YFR057W* Ct on each sample and then a ∆∆Ct was calculated for each replicate as described in (Schmittgen and Livak 2008; Livak and Schmittgen 2001) relative to WT strain BY4741; average fold-change expression and SD were calculated.

## ACKNOWLEDGEMENTS

We thank members of the DeLuna lab for helpful discussions. We are grateful to Irene Castaño for useful suggestions and critical reading of the manuscript and to Cristina Aranda for technical assistance. This work was funded by the Consejo Nacional de Ciencia y Tecnología de México (CONACyT grants CB-2015/164889, FORDECYT-PRONACES/103000/2020, and CB-2016-01/282511). AJ-R was supported by a CONACyT postdoctoral fellowship (167877), funding from Cinvestav and a Welcome Trust Seed Award in Science to EM. The funders had no role in study design, data collection and analysis, decision to publish, or preparation of the manuscript.

## DATA AVAILABILITY

Strains and plasmids are available upon request. All datasets generated for this study are included in the article or Supplementary Materials.

## SUPPLEMENTARY MATERIAL

**Supplementary Note S1**. List of genes previously associated with telomeric silencing in *Saccharomyces cerevisiae*.

**Supplementary Table S1**. Features of different subtelomeric loci and silencing analysis.

**Supplementary Table S2**. Potentially novel telomeric silencing genes in *Saccharomyces cerevisiae*

**Supplementary Table S3**. List of genes used for *kappa* statistical analysis.

**Supplementary Table S4**. List of strains used in this study.

**Supplementary Table S5**. List of primers used in this study.

**Supplementary Figure S1**. Insertion of the *URA3-*GFP does not disrupt nucleosome positioning at the *YFR057W* promoter.

**Supplementary Figure S2**. *Si score* of two independent screens at the *COS12* locus are correlated.

**Supplementary Figure S3**. Validation of *Si scores* of selected mutants and analysis of the silencing level by relevant groups.

**Supplementary Dataset S1**. Raw data of subtelomeric silencing screens *COS12* and *YFR057W* by flow cytometry (XLS).

